# Robust control of replication initiation in the absence of DnaA-ATP ↔ DnaA-ADP regulatory elements in *Escherichia coli*

**DOI:** 10.1101/2022.09.08.507175

**Authors:** Thias Boesen, Godefroid Charbon, Haochen Fu, Cara Jensen, Michael Sandler, Suckjoon Jun, Anders Lobner-Olesen

## Abstract

Investigating a long-standing conceptual question in bacterial physiology, we examine why DnaA, the bacterial master replication initiator protein, exists in both ATP and ADP forms, despite only the ATP form being essential for initiation. We engineered the Δ4 *Escherichia coli* strain, devoid of all known external elements facilitating the DnaA-ATP/ADP conversion, and found that these cells display nearly wild-type behaviors under non-overlapping replication cycles. However, during rapid growth with overlapping cycles, Δ4 cells exhibit initiation instability. This aligns with our model predictions, suggesting that the intrinsic ATPase activity of DnaA alone is sufficient for robust initiation control in *E. coli* and the DnaA-ATP/ADP conversion regulatory elements extend the robustness to multifork replication, indicating an evolutionary adaptation. Moreover, our experiments revealed constant DnaA concentrations during steady-state cell elongation in both wild-type and Δ4 cells. These insights not only advance our understanding of bacterial cell-cycle regulation and DnaA, but also highlight a fundamental divergence from eukaryotic cell-cycle controls, emphasizing protein copy-number sensing in bacteria versus programmed protein concentration oscillations in eukaryotes.

## Introduction

Initiation of chromosome replication in bacteria is a highly regulated process, intricately linked with cellular growth and division. This precision is evident in the remarkably consistent cell size at initiation observed in *Escherichia coli* and *Bacillus subtilis*, with a coefficient of variation (CV) of no more than 10% - notably lower than other physiological variables with CVs around 20% (Si et al., 2019; Sauls et al., 2019a; Wallden et al., 2016; Knöppel et al., 2023; Si et al., 2017). These bacteria maintain a stable initiation mass (the cell size per origin) across various growth conditions, exemplified by *E. coli*’s initiation mass varying by a maximum of 20% over a 10-fold growth rate range (Si et al., 2017; Donachie, 1968; Wold et al., 1994; Zheng et al., 2020).

The molecular biology of initiation in *E. coli* has been extensively studied, particularly focusing on its master regulator, DnaA. Replication begins at the *oriC* origin, initiated by DnaA binding. DnaA functions as an AAA+ ATPase, existing in either an active (DnaA-ATP) or inactive (DnaA-ADP) form. The *oriC* region contains both high- and low-affinity binding sites for the two forms of DnaA. Though the high-binding sites have similar binding affinities to both DnaA-ATP and DnaA-ADP, only DnaA-ATP can bind the low-binding sites helped by cooperativity (McGarry et al., 2004; Ozaki et al., 2008). The binding of DnaA-ATP at these sites leads to oligomerization and helical filament formation, generating tension that opens the duplex at the adjacent DUE site, and facilitating replisome assembly (Sekimizu et al., 1987; Erzberger et al., 2006; Leonard and Grimwade, 2011; Riber et al., 2016). This mechanism underscores the necessity of DnaA-ATP for initiation.

Several factors influencing the DnaA-ADP ↔ DnaA-ATP conversion have been identified. DARS1 and DARS2 (<u>D</u>na<u>A r</u>ejuvenating <u>s</u>equences 1 and 2) facilitate the conversion from DnaA-ADP to DnaA-ATP, while RIDA (<u>r</u>egulatory <u>i</u>nactivation of <u>D</u>na<u>A</u>) and DDAH (*<u>d</u>atA*-dependent DnaA-ATP <u>h</u>ydrolysis) mechanisms promote the reverse conversion (Kato and Katayama, 2001; Fujimitsu et al., 2009; Kasho and Katayama, 2013). These elements’ perturbations impact initiation precision and synchrony to varying degrees (Frimodt-Møller et al., 2016; Kitagawa et al., 1998; Charbon et al., 2014).

Despite these discoveries, the role of DnaA-ADP in *E. coli*, when only DnaA-ATP is needed for initiation, remains unclear. It is hypothesized that the DnaA-ADP ↔ DnaA-ATP conversion creates periodic oscillations in their concentrations, with replication initiating when the ratio of [DnaA-ATP]/[DnaA-ADP] peaks, akin to eukaryotic cell-cycle control (Donachie and Blakely, 2003; Berger and Wolde, 2022; Fu et al., 2023). However, this oscillation theory contrasts with the concept of balanced biosynthesis in bacterial physiology, where key division proteins like FtsZ maintain a near-constant concentration during cell elongation (Jun et al., 2018). We previously proposed, following the initiator titration model (Hansen et al., 1991), that DnaA-ADP is unnecessary for initiation but helps stabilize it by preventing re-initiations (Fu et al., 2023).

In this study, we show that DnaA’s intrinsic ATPase activity alone is sufficient for precise and robust initiation in *E. coli*. We created the “Δ4” *E. coli* strain, devoid of all known external DnaA-ADP ↔ DnaA-ATP conversion elements. This strain not only survives but also exhibits near wild-type behavior under non-overlapping replication cycles. However, under richer nutrient conditions that lead to overlapping replication cycles, replication re-initiations occur, aligning with model predictions (Berger and Wolde, 2022; Fu et al., 2023). Additionally, our single-cell level analysis reveals balanced DnaA synthesis, despite dnaA promoter’s weak auto-repression or sequestration. These findings offer new insights into bacterial cell cycle control, its evolutionary aspects, and fundamental distinctions from eukaryotic mechanisms.

## Results

### DnaA-ATP ↔ DnaA-ADP conversion and construction of the Δ2^ATP^ and Δ2^ADP^ E. coli cells

Currently, four extrinsic factors for the DnaA-ADP ↔ DnaA-ATP conversion are known (Figure 1.1). For DnaA-ADP → DnaA-ATP, DARS1 and DARS2 at different loci on the chromosome have been characterized (Fujimitsu et al., 2009; Kasho et al., 2014; Sugiyama et al., 2019). For DnaA-ATP → DnaA-ADP, the two extrinsic mechanisms are RIDA mediated by Hda and the DNA-loaded *β-*sliding clamps (Kato and Katayama, 2001; Nishida et al., 2002; Su’etsugu et al., 2005, 2004, 2013), and DDAH mediated at the five DnaA boxes at the datA locus (Kitagawa et al., 1998; Kasho and Katayama, 2013).

**Figure 1.**
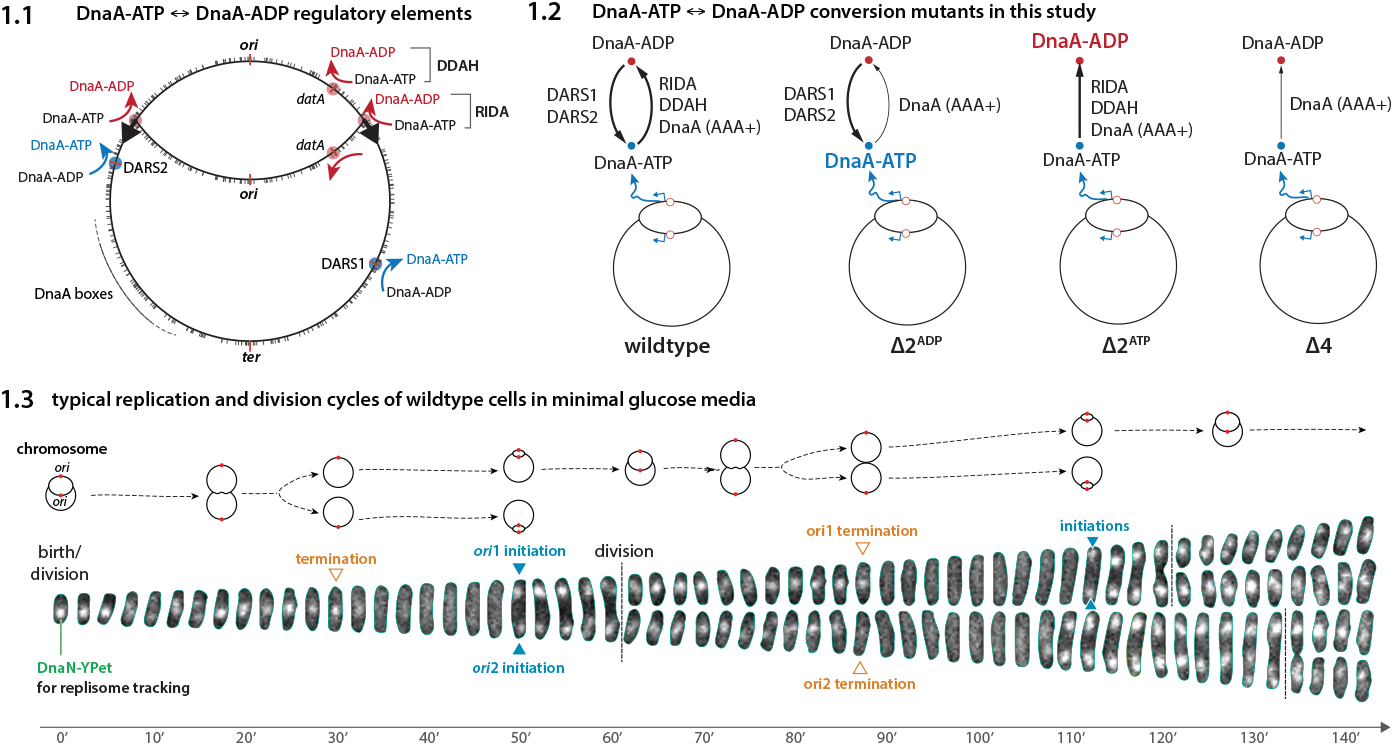
Extrinsic DnaA-ATP ↔ DnaA-ADP conversion elements. **(1.1)** The four extrinsic DnaA-ATP ↔ DnaA-ADP conversion regulatory elements. **(1.2)** We constructed Δ2^ATP^ (ΔDARS1 ΔDARS2), Δ2^ADP^ (Δ*hda* Δ*datA*), and Δ4 (ΔDARS1 ΔDARS2 Δ*hda* Δ*datA*) *E. coli* strains. **(1.3)** Tracking the replication dynamics using fluorescence-labeled replisomes (DnaN-YPet) (Si et al., 2019) (see Materials and Methods). Chromosome configurations (tracking only the top cells) over time are shown above. The wild-type cells typically initiate synchronously.

To examine the impact of the extrinsic DnaA-ADP → DnaA-ATP vs. DnaA-ATP → DnaA-ADP regulatory elements on initiation, we constructed two types of *E. coli* strains: Δ2^ATP^ (ΔDARS1 and ΔDARS2) and Δ2^ADP^ (Δ*hda* + Δ*datA*) strains. Therefore, in the Δ2^ATP^ cells, the only source of DnaA-ATP is the de novo synthesis of DnaA, which becomes DnaA-ATP due to the high cytoplasmic ATP concentration. The Δ2^ATP^ cells have lower [DnaA-ATP] and DnaA-ATP/DnaA-ADP ratio than the wild-type cells (Riber et al., 2016; Fujimitsu et al., 2009).

By contrast, the Δ2^ADP^ cells must exclusively rely on the *intrinsic* ATPase activity of DnaA to convert DnaA-ATP to DnaA-ADP. Consequently, the Δ2^ADP^ cells exhibit a higher level of DnaA-ATP and the DnaA-ATP/DnaA-ADP ratio (Kasho and Katayama, 2013).

### DnaA-ATP →DnaA-ADP and DnaA-ADP → DnaA-ATP have opposing effects on initiation

One of the main goals of the present study is to differentiate the impact of initiation mutants on initiation stability and the role of DnaA-ATP ↔ DnaA-ADP conversion. For example, wild-type cells generally show highly synchronized initiation (Figure 1.3), even when the initiation mass can vary significantly from cell to cell.

The single-cell initiation data show stark differences between Δ2^ATP^ vs. Δ2^ADP^ cells (Figure 2). Firstly, Δ2^ATP^ exhibits a notable initiation delay. The initiation mass distribution is shifted to the right, with an approximately 70% larger average and a slightly increased CV than the wildtype (Figure 2.1). Flow cytometry data indicates that most cells contain either one or two replication origins (ori’s) (Figure 2.1, Figure S6), suggesting non-overlapping cell cycles. In other words, initiation typically occurs after cell birth, a trend supported by the fork plot in Figure 2.3 (see Materials and Methods) and illustrated in a real example in Figure 2.2. Noticeably, the replisome foci distribution in the cell is highly symmetric in this mutant, which differs from the discovery in a previous study of a mutant with a similar genotype (Knöppel et al., 2023).

**Figure 2.**
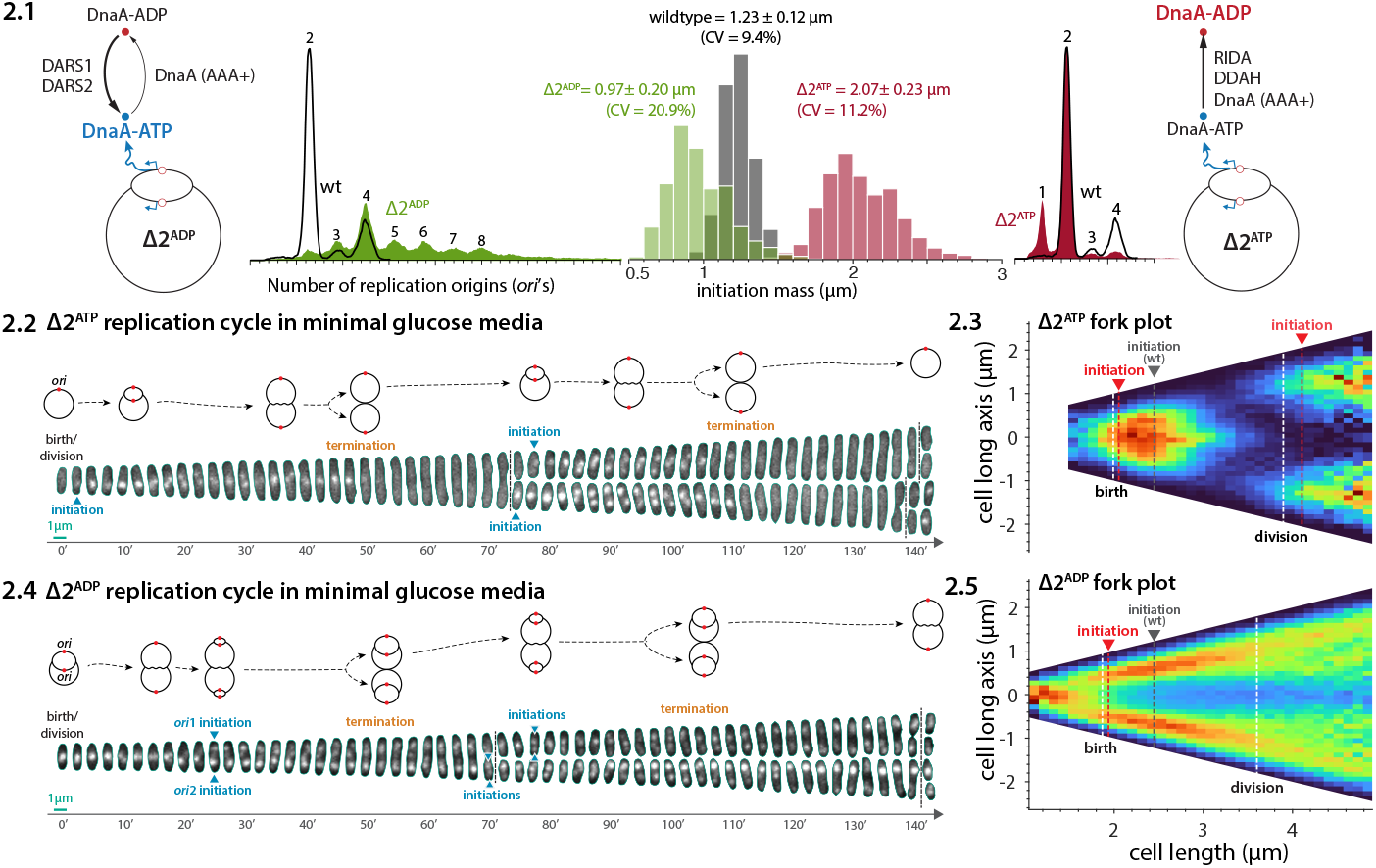
Opposing effects of the Δ2^ATP^ and Δ2^ADP^ mutants on initiation. **(2.1)** The initiation mass distributions from mother machine experiments and the flow cytometry data. **(2.2, 2.4)** Timelapse fluorescence imaging of Δ2^ATP^ (Δ2^ADP^) cells with chromosome configurations above. **(2.3, 2.5)** The fork plot of the Δ2^ATP^ (Δ2^ADP^) cells shows the distribution of foci along the cell long axis in cells binned by cell length. The dashed mean initiation lines are from the initiation mass dstributions from (A).

**Figure 3.**
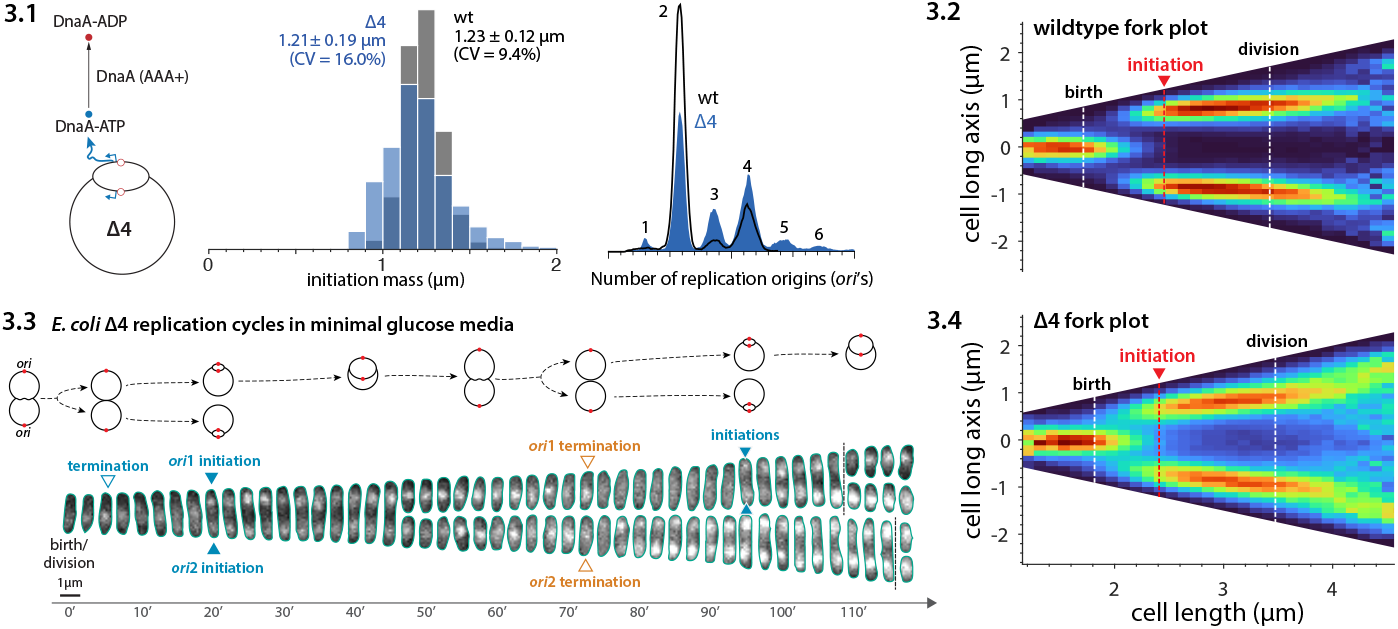
Robustness of Δ4 *E. coli*. **(3.1)** The initiation mass distributions and the flow cytometry data for Δ4 cells vs. wildtype cells. **(3.2, 3.4)** fork plots for Δ4 cells vs. wildtype cells. **(3.3)** Timelapse fluorescence imaging of Δ4 cells with chromosome configurations shown above.

By contrast, Δ2^ADP^ initiates prematurely, as evidenced by the smaller initiation mass than the wildtype cells (Figure 2.1). The average initiation mass is smaller by around 20%, while the CV increases significantly from 10% to above 20% (Figure 2.1). This mutant was particularly difficult to analyze due to its highly noisy initiation behavior. For example, unlike the wild-type cells in the same growth condition, Δ2^ADP^ cells typically undergo overlapping cell cycles with a new round of replication initiates before the previous round of replication terminates (Figure 2.4), a challenging condition for tracking replication cycles using image analysis (see Materials and Methods). Based on the fork plot in Figure 2.5, we suspect that the actual mean initiation mass could be even smaller and the CV larger than our statistics suggest (see Figure S5 for estimation details).

**Figure 4.**
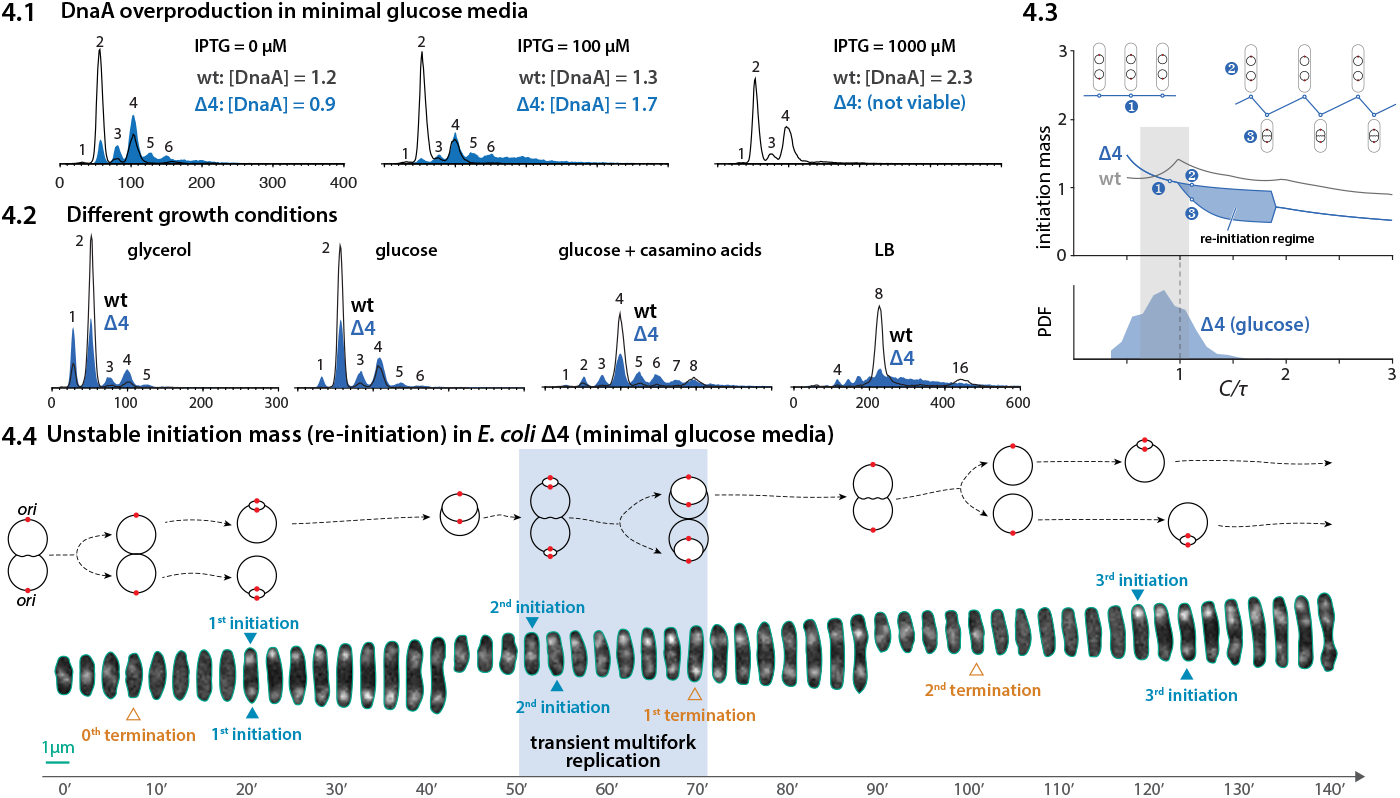
Δ4 cells vs. wildtype cells in perturbations of DnaA concentration and growth conditions. **(4.1)** Flow cytometry data of wildtype and Δ4 cells under different induction level of an extra copy of *dnaA* gene on a plasmid. **(4.2)** Flow cytometry data of wildtype and Δ4 cells under different growth conditions. **(4.3)** Theoretical predictions from the initiator-titration model v2. Lower panel: The distribution of *C/τ* for Δ4 cells in the glucose condition. **(4.4)** A time-lapse imaging of 4 cells in the glucose condition as an example of unstable initiation mass. The chromosome configurations over time are illustrated above.

**Figure 5.**
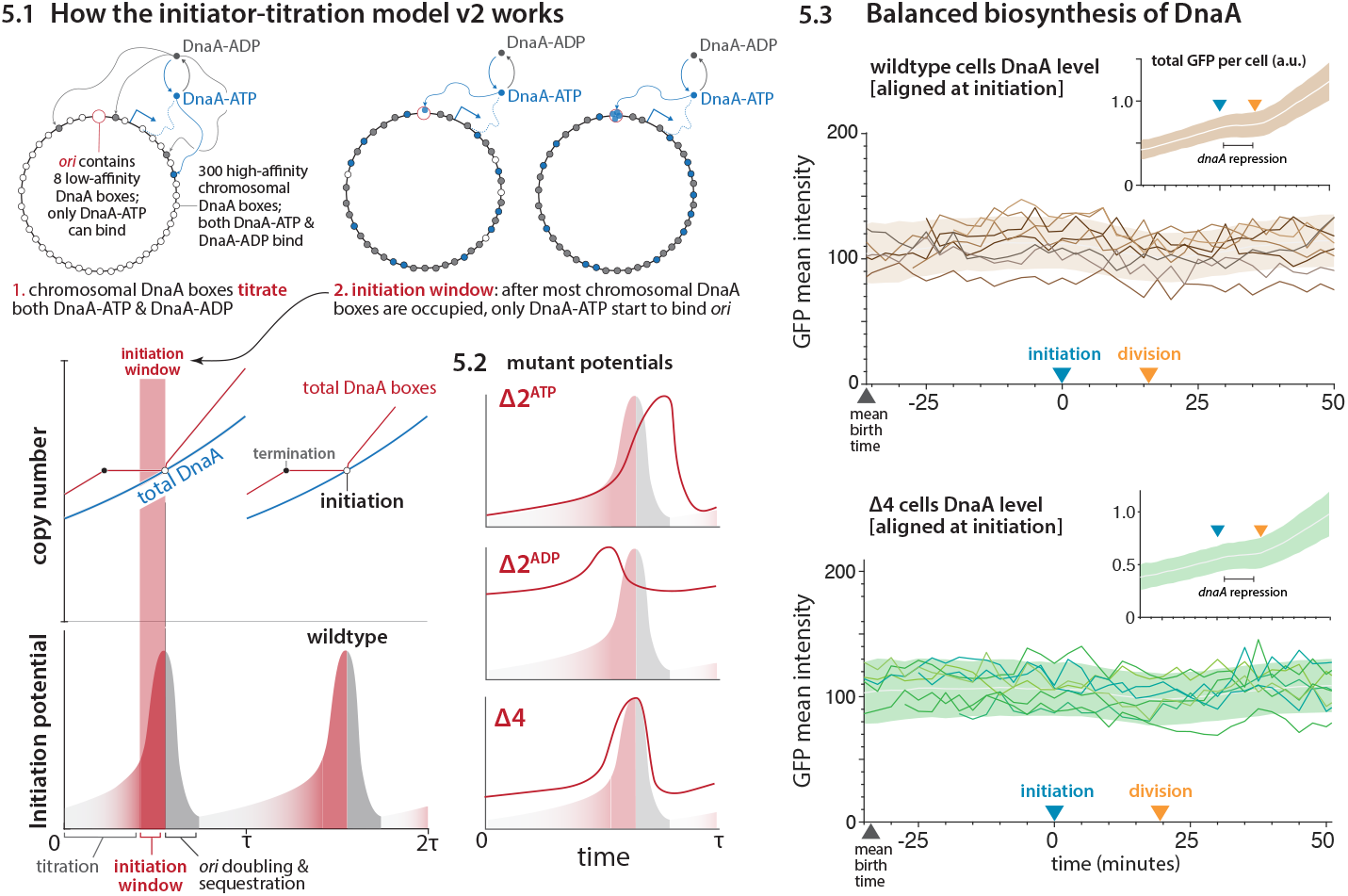
The initiator-titration model v2 and the Δ2/Δ4 mutant behaviors. **(5.1)** The initiator-titration model v2 and the initiation potential. **(5.2) Δ**2 and Δ4 initiation potential. **(5.3)** The concentration of DnaA is nearly constant in both wildtype and Δ4 E. coli. Therefore, [DnaA-ATP] and [DnaA-ADP] also must be constant in Δ4 E. coli during steady-state cell elongation.

These results indicate that the DnaA-ATP → DnaA-ADP and DnaA-ADP → DnaA-ATP conversion elements have opposing effects on initiation.

### Δ4 E. coli exhibits a near wild-type phenotype despite the absence of all known extrinsic DnaA-ATP DnaA-ADP conversion elements

As the Δ2^ATP^ (ΔDARS1 ΔDARS2) and Δ2^ADP^ (Δ*hda* Δ*datA*) show the opposing effects, we wondered if we could delete all four extrinsic conversion elements (DARS1, DARS2, *hda*, and *datA*) (Figure 3.1). To the best of our knowledge, such attempts had never been made before, probably because of the apparent importance of DnaA-ADP ↔ DnaA-ATP in initiation control. Indeed, deletion of each of these elements typically impacts the cells adversely, as seen in Figure 2 (Kato and Katayama, 2001; Charbon et al., 2014, 2018).

To our surprise, however, the Δ4 (ΔDARS1 ΔDARS2 Δ*hda* Δ*datA*) *E. coli* not only was viable but also exhibited a near wild-type initiation phenotype. Specifically, the average initiation mass remained similar to the wild-type cells, with a modest increase in the CV of the initiation mass from 9.4% to 16.0% (Figure 3.1). This is consistent with the flow-cytometry data, which shows a small peak for three chromosomes due to the slightly increased initiation asynchrony (Figure 3.1, Figure S6). The statistics and flow cytometry data are also consistent with the fork plots (Figs. 3.2 and 3.4), where Δ4 and wildtype cells show similar fork patterns but Δ4 is noisier. Further, the Δ4 and wildtype have similar average values in other parameters, including the growth rate and cell sizes, although the CV is generally larger in Δ4 (Figure S1).

These results suggest that the rate of DnaA-ADP → DnaA-ATP by DARS1 + DARS2 and DnaA-ATP → DnaA-ADP by *hda* + *datA* (RIDA and DDAH) must be comparable to each other in the wild-type cells. In the Δ4 cells, their opposing effects should largely cancel out, only mildly compromising the synchrony of the initiation processes.

### The intrinsic ATPase activity of DnaA can be sufficient for the wild-type phenotype

The unexpected robust initiation control in the Δ4 *E. coli* raises the fundamental question: if only DnaA-ATP is required for initiation at ori, why does *E. coli* maintain both forms of DnaA and the intricate DnaA-ATP → DnaA-ADP conversion elements? We present two orthogonal experimental results that show both forms of DnaA are required to improve initiation stability to a wider range of physiological conditions.

First, we gradually overproduced wild-type DnaA (DnaA^WT^) in the Δ4 background in intermediate growth conditions (Figure 4.1). The Δ4 cells were viable up to 1.7-fold the wild-type level of DnaA, and showed re-initiation beyond that point. By contrast, wild-type cells tolerated a significantly higher level of DnaA and only over-initiated modestly (Figure 4.1). Nevertheless, that the Δ4 *E. coli* is viable at the elevated level of DnaA suggests a significant ATPase activity of DnaA *in vivo*.

Next, we also investigated Δ4 *E. coli* ‘s tolerance of two different DnaA mutants. DnaA^R334A^ is known for a significantly decreased ATPase activity (thus a higher DnaA-ATP level) (Nishida et al., 2002; Nakamura and Katayama, 2010), whereas DnaA^T174P^ shows an increased activity (thus a lower DnaA-ATP level) (Gon et al., 2006). The Δ4 cells did not tolerate additional DnaA^R334A^ at all, likely due to the elevated DnaA-ATP level, compared to the wild-type cells (Figure S2.1). Flow-cytometry data indicated over-initiation with a varying degree in both Δ4 and wild-type cells with DnaA^R334A^ (Figure S2.1).

On the other hand, the fast ATPase DnaA^T174P^ variant did not cause a significant change in the average initiation mass at low to intermediate overproduction for both Δ4 and wild-type cells. Upon strong overproduction of DnaA^T174P^, initiation was delayed (Figure S2.2). We interpret these results as DnaA-ATP → DnaA-ADP in Δ4 was already sufficient for stable initiation, until high levels of DnaA-ADP by DnaA^T174P^ became detrimental to the initiation process. One possible mechanistic cause is the formation of short DnaA-ADP oligomers following the de novo synthesis of DnaA^T174P^, making DnaA-ATP unable to compete for initiation in this strain (Kawakami et al., 2005).

These results underscore that the intrinsic ATPase activity is sufficient for the observed wild-type phenotype of Δ4 in intermediate growth conditions, but their ability to buffer the impact of DnaA-ATP or DnaA-ADP level perturbations is less than the wild-type cells.

### Extrinsic DnaA-ATP ↔ DnaA-ADP conversion elements improve initiation stability to multifork replication

To assess the robustness of Δ4 cells to physiological perturbations, we tracked the replication cell cycle in various growth conditions. Interestingly, Δ4 cells exhibit normal growth and replication initiation under slow growth conditions (Figure 4.2, Figure S4). However, their differences become significant under fast growth conditions, with the Δ4 *E. coli* showing more asynchronous initiations (Figure 4.2), resulting in growth defects (Figure S4 & S5). These results suggest that Δ4 cells are less robust against perturbations in fast growth conditions compared to wild-type cells.

Previously, we predicted using our initiator-titration model v2 that *E. coli* cells lacking external DnaA-ATP ↔ DnaA-ADP conversion elements can develop initiation instability during multifork replication (Fu et al., 2023). In this model, the initiation mass can oscillate between two values as the mass-doubling time becomes shorter than the C period (*C/τ >* 1) (Figure 4.3). This unstable initiation can span a significant range of physiological conditions, shown as the shaded island in the *C/τ* vs. initiation mass in Figure 4.3. When two consecutive initiation events are close enough to be in the same generation, re-initiation occurs.

In the glucose minimal medium, *C/τ* was 0.84 ± 0.25 for Δ4 *E. coli* (Figure 4.3). Therefore, a subpopulation of Δ4 cells can enter the multifork replication regime (*C/τ >* 1) due to the large variation of the doubling time and C period. Indeed, such predicted oscillations can be seen in our single-cell tracking experiment with *E. coli* Δ4 (Figure 4.4).

In rich media (MOPS rich glycerol), we measured the average *C/τ ≈* 1.2 − 1.3 by mother machine (Figure S5), indicating that most Δ4 cells are in the re-initiation regime (Figure 4.3). We were unable to track the replication cycles at the single-cell level due to significant re-initiations and asynchrony. Nevertheless, the noisy initiation in Δ4 *E. coli* observed in both flow-cytometry (peaks at the odd numbers) and mother machine experiments is well-aligned with the predicted initiation instability.

The initiation instability of Δ4 *E. coli* during multifork replication can be understood intuitively. Δ4 *E. coli* lacks, among others, DnaA-ATP → DnaA-ADP conversion at the replication fork by RIDA. Therefore, Δ4 *E. coli* has a higher level of DnaA-ATP compared to wild-type *E. coli*, making Δ4 *E. coli* cells prone to re-initiation (see the next section and Figure 5). Based on these results, we suggest that the role of extrinsic DnaA-ATP ↔ DnaA-ADP conversion elements is to extend initiation stability to multifork replication.

### The initiator-titration model v2 can explain the Δ2 and Δ4 behaviors

Recently, we extended the initiator-titration model originally proposed by Hansen and colleagues (Fu et al., 2023; Hansen et al., 1991). In this initiator-titration model v2, we took into account the conversion of DnaA-ATP and DnaA-ADP. The key idea underlying both versions of the model is that DnaA (either form) proteins are first titrated by the DnaA boxes distributed along the chromosome before initiation, dividing the replication cycle into two sequential stages, namely, titration followed by initiation (Figure 5.1).

During the titration stage, the initiation “potential” is strongly suppressed. As the number of unoccupied chromosomal DnaA boxes becomes comparable to the DnaA boxes within *oriC*, DnaA-ATP molecules will start to bind *oriC*. Upon initiation, the DnaA boxes within *oriC* double in number, and initiation is sequestered by SeqA (Lu et al., 1994). The initiation potential thus drops significantly upon initiation and remains low during this period due to RIDA and DDAH activity during sequestration.

By translating the initiator-titration model v2 to initiation potential, we can understand the behavior of Δ2^ATP^, Δ2^ADP^, and Δ4 in the following manner (Figure 5.2).

1. Δ2^ATP^: the level of DnaA-ATP is lower than the wildtype, thus initiation delays since binding of the lowered level of DnaA-ATP to *oriC* is slowed down. DnaA titration and the related initiation potential are largely unaffected since both DnaA-ATP and DnaA-ADP can bind the high-affinity chromosomal DnaA boxes.
2. Δ2^ADP^: The overall potential is markedly elevated due to the significantly higher level of DnaA-ATP, and it remains high after initiation due to the absence of RIDA and DDAH, both of which provide negative feedback on initiation (Fu et al., 2023). This can cause re-initiations.
3. Δ4: The overall initiation potential is still elevated due to the lack of RIDA and DDAH. Δ4 *E. coli* also lacks DARS1/2 and thus has a reduced DnaA-ADP → DnaA-ATP activity, alleviating the effect of missing RIDA and DDAH. However, due to the generally elevated level of DnaA-ATP, a modest additional increase in DnaA-ATP can cause re-initiation in the Δ4 cells, as seen in the DnaA^R334A^ mutant experiment discussed earlier (Figure S2.1).

### Both wildtype and Δ4 E. coli show nearly constant concentrations of DnaA during cell elongation

One of the major assumptions of the initiator-titration models is the balanced biosynthesis of DnaA. In the practical aspect, balanced biosynthesis leads to constant protein concentration during cell elongation. This is in stark contrast to eukaryotic cell cycle controls dominated by programmed gene expression and protein degradation, causing changes in protein concentrations, such as oscillations for cyclins.

We tracked the DnaA level using a fast-maturing GFP in the mother machine (Wang et al., 2010). We found that the DnaA level is indeed practically constant during steady-state cell elongation (Figure 5.3). However, this is at odds with the notion of *dnaA* autoregulation or *dnaA* repression by SeqA, which should decrease DnaA concentrations (Lu et al., 1994; Campbell and Kleckner, 1990; Theisen et al., 1993; Løbner-Olesen et al., 2003). Intriguingly, careful examination of the total amount of DnaA (by integrating the GFP signal across the cell) vs. time revealed a pause in DnaA production (Figure 5.3 insets), consistent with periodic repression of *dnaA* expression after initiation. Nevertheless, as seen in the DnaA concentration vs. time, the impact of *dnaA* repression is modest compared to the scale of protein concentration fluctuations (Figure 5.3). We thus conclude that DnaA production is nearly balanced, well-aligned with the initiator-titration models (Fu et al., 2023; Hansen et al., 1991) and the adder principle of size control (Si et al., 2019; Sompayrac and Maaløe, 1973).

The constancy of DnaA level in Δ4 *E. coli* has important implications. To see this, consider the balanced biosynthesis of DnaA, 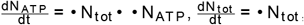, and 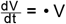, where *N*_ATP_ is the number of DnaA-ATP, *N*_tot_ the total number of DnaA, *V* the cell volume, and λ the growth rate. The steady-state solution of these equations leads to [DnaA] = constant (Figure 5.3), [DnaA-ATP] = constant, and [DnaA-ADP] = constant during cell elongation. Since the protein concentrations are constant, the cell must accumulate a threshold number of initiator proteins to trigger a cell cycle event, such as DnaA for replication initiation (Si et al., 2019; Sompayrac and Maaløe, 1973) or FtsZ for cell division (Si et al., 2019). These conclusions represent a fundamental difference from eukaryotic cell cycle control based on oscillations of cyclin concentrations via programmed gene expression and/or protein degradation (Hunt and Murray, 1993).

## Discussion

In this work, we have shown that Δ4 *E. coli*, devoid of all known extrinsic regulatory elements for DnaA-ATP ↔ DnaA-ADP conversion (DARS1, DARS2, *hda*, and *datA*), can exhibit near wild-type phenotype for initiation control. A new emerging picture from this study and our previous works (Si et al., 2019; Fu et al., 2023) is that the bacterial cell cycle control must depend on the protein counting mechanisms, rather than protein concentration sensing in eukaryotes. Since balanced biosynthesis is the hallmark of bacterial physiology (Jun et al., 2018), the differences between bacteria and eukaryotes are profound and merit further investigations in the future.

Apart from initiation, our Δ2 and Δ4 mutants show interesting features that warrant future investigation. For example, while Δ4 and Δ2^ADP^ cells show co-localized replisomes (Figs. 1.3, 3.3, and 2.4) consistent with recent reports (Mangiameli et al., 2018; Gras et al., 2023; Youngren et al., 2014), Δ2^ATP^ cells show significantly different behaviors with one replisome focus splitting into two foci during DNA replication (Figure 2.2). It remains to be seen whether these differences reflect distinct overall chromosome organization as suggested previously (Youngren et al., 2014) or more active interactions between replication forks.

Furthermore, more insightful information may be within reach based on new high-throughput data and mathematical modeling (Huang et al., 2023; Pflug et al., 2023; Bhat et al., 2023; Le Treut et al., 2021), which can reveal the relationship between initiation and replication dynamics.

Finally, the robust initiation of the Δ4 cells and its mechanistic implications was a major surprise for us. At the same time, we also find them gratifying from an evolutionary perspective (Pelletier et al., 2021; Olivi et al., 2021).

For example, the First Cell would only need one master regulator protein with intrinsic ATPase activity, and different organisms may have acquired organism-specific additional control mechanisms to extend the physiological space for robust initiation control and stability during evolution (Figure 4). While modest in the cell-cycle timescale, their impact on the evolutionary timescale is unquestionable.

## Methods

### Growth conditions

In flow cytometry experiments and most of the mother machine experiments, cells were grown in Lysogeny Broth (LB) or AB minimal medium (Clark & Maaløe, 1967) supplemented with 10 *μ*g/mL thiamine and either 0.2% glucose (ABT-Glycerol) or 0.2% glycerol (ABT-Glucose). Only in the mother machine experiments in fast growth conditions (Figure S5), we used MOPS rich media (Si et al., 2017; Neidhardt et al., 1974) supplemented with 0.2% glycerol. When necessary, antibiotics were added in the following concentrations: kanamycin, 50 *μ*g/mL; chloramphenicol, 20 *μ*g/mL; ampicillin, 150 *μ*g/mL.

### Construction of mutants of DnaA-ATP ↔ DnaA-ADP regulatory elements

All strains used were derivatives of E. coli K-12 MG1655 (F-λ-rph-1) (Guyer et al., 1981). All deletions were performed by P1-mediated transduction (Miller, 1972). The Δ*hda* Δ*datA* strain was constructed by transducing Δ*hda* into a Δ*datA* strain and plating it on an ABT-Glycerol medium as described previously (Charbon et al., 2018; Riber et al., 2006a). The individual DARS1:*cat* (ALO4313) and DARS2:*cat* (ALO4254) mutants were constructed previously (Frimodt-Møller et al., 2016, 2015) using the λ-red procedure (Datsenko and Wanner, 2000). Before combining the ΔDARS1 and ΔDARS2, the *cat* resistance marker was cured from ALO4313 using pCP20 (Cherepanov and Wackernagel, 1995). The Δ4 strain (ΔDARS1 ΔDARS2 Δ*hda* and Δ*datA*) was constructed by combining the above by P1 transduction. All strains were whole-genome sequenced to confirm the presence of the mutations and to ensure that no suppressors appeared (see SI Table S2).

### Construction of fluorescence-labeled strains

To track the initiation of chromosome replication in the strains, we inserted a fluorescence-labeled sliding-clamp of the replisome, *dnaN-yPet* (Si et al., 2019; Reyes-Lamothe et al., 2010), into the wildtype, Δ2^ATP^, Δ2^ADP^, and Δ4 strains through P1 transduction (Table S1).

To measure the DnaA level while tracking initiations in the wildtype and Δ4, we inserted a fast-maturing GFP protein, *gfpmut* 2 (Balleza et al., 2018), between *dnaA* and *dnaN* as a transcription reporter for *dnaA*. Due to the similarity of the emission spectra between YPet and GFPmut2, we used mCherry instead of YPet as a fluorescenct tag for the replisome. The *gfpmut* 2 and the *mcherry* genes were assembled with an ampicillin resistance gene using Gibson assembly, and the entire fragment was inserted to the *dnaA* operon on the chromosome of both the wildtype and Δ4 through recombineering (Thomason et al., 2014). For Δ4, we verified the loci of previous deletions through sequencing after recombineering (Table S1).

### Plasmids

Overexpression of DnaA was performed using pLR40 (Riber et al., 2006b).

The pLRR334A with DnaA^R334A^ was created using the primers: CTG CAG GTC GAC GGA TCC CCA A, AGC CCA GCG CGT CGG CCG CCA T, TCGCCAAGCGTCTACGATCTAACGTAGCTGAGCTGGAAGGGGCGCTGAAC and GTTCAGCGCCCCTTCCAGCTCAGCTACGTTAGATCGTAGACGCTTGGCGA.

The pLRT174P with DnaA^T174P^ was created using primers: CTG CAG GTC GAC GGA TCC CCA A, AGC CCA GCG CGT CGG CCG CCA T, ATGGCGGCCCGGGTCTGGGTAAAACTCACC and ACCCAGACCCGGGCCGCCATAAAGGAACAA.

### DnaA level measurements

Samples for western blots were prepared as described previously (Charbon et al., 2021). Quantification of protein levels and analysis was performed using ImageJ software and the signal relative to the wildtype (ALO8238) was calculated.

### Flow cytometry

Cells were balanced in exponential growth for more than 10 generations and incubated with 300 *μ*g/mL rifampicin and 36 *μ*g/mL at 37°C for 4 hours, allowing ongoing replication cycles to terminate. Cells were harvested and resuspended in 100 *μ*L 10 mM Tris-HCl, pH 7.4, and fixed by adding 1 mL of 77% ethanol and stored at 4°C. Before flow cytometric analysis, fixed cells were pelleted by centrifugation at 15,000 rcf for 15 min. The supernatant was discarded, and the pellet was resuspended in 150 *μ*l staining solution (90 *μ*g/mL mithramycin, 20 *μ*g/mL ethidium bromide, 10 mM MgCl2, 10 mM Tris-HCl, pH 7.4). Samples were kept on ice for a minimum of 10 minutes before analysis.

We performed flow cytometry experiments as described previously (Løbner-Olesen et al., 1989) using an Apogee A10 system. A minimum of 30,000 cells for each sample were analyzed, and the number of origins per cell and relative mass were determined as described previously (Løbner-Olesen et al., 1989).

### Microfluidic experiments

We used PDMS-based mother machines to track single-cell lineages (Si et al., 2019; Taheri-Araghi et al., 2015; Wang et al., 2010). To guarantee the same experimental conditions for control experiments, we designed duplex mother machines and multiplex mother machines that contain two main trenches and sixteen main trenches on each device, respectively (Figure S7). Each side of the main trench consists of 2000 channels with a width ranging from 1.1 *μ*m to 1.5 *μ*m in design. The inlet of each main trench was connected to a syringe with a growth medium by Teflon tubing. The syringes were controlled by syringe pumps (PHD Ultra, Harvard Apparatus, MA) to maintain a constant flow rate.

In preparation of a mother-machine experiment, a single colony was inoculated in 1 mL ABT-Glucose, containing appropriate antibiotics, and incubated overnight at 37°C in a water-bath shaker. Cells were then diluted 1000-fold into fresh media and balanced in an exponential phase for more than 10 generations. When reaching an OD^600^ of 0.1-0.15, cells were pelleted, concentrated, and injected into the mother machine device prewashed with BSA to reduce cell adhesion in the device channels. A constant flow of fresh growth media was supplied through the device for the duration of the experiment, as described previously (Si et al., 2019; Taheri-Araghi et al., 2015). After setting up the microfluidics and the microscopy program, we waited around 3 hours before imaging to ensure cells were in the steady state.

### Microscopy

We performed phase contrast and fluorescence imaging tandemly on an inverted microscope (Nikon Ti-E) with Perfect Focus 3 (PFS3), 100x oil immersion objective (PH3, numerical aperture = 1.45), Obis lasers 488LX (Coherent Inc., CA) as fluorescence light source, Chroma EYFP filter (49003-ET-EYFP), and Prime 95B sCMOS camera (Photometrics) (Si et al., 2019). The laser power for 488 nm excitation was 30 mW, and the exposure time was 100 ms. The imaging time interval was 2.5 min for all the mother machine experiments.

### Image analysis

We used our own Napari-mm3 (Thiermann et al., 2023), following the developed protocol of mother machine image analysis (Thiermann et al., 2023; Sauls et al., 2019b).

For cell segmentation, we used a machine-learning-based method, U-net, rather than the traditional Otsu method to obtain more reliable results (Ronneberger et al., 2015).

The cell segmentation results were also used in plotting time-lapse fluorescence images, such as in Figure 1.3. In Adobe Illustrator (2023), a segmentation image and the corresponding fluorescence image are overlapped first, and the “image trace” function is applied to the segmentation image to track and smoothen the boundary of each segmented cell. We then used the function “make clipping mask” to obtain individual cell images with fluorescence inside.

For fork plots, we used a Laplacian of Gaussian method to identify fluorescence foci, as described previously (Si et al., 2019). The position of foci along the cell’s long axis (y-axis) is recorded and binned by 1 pixel of the original image (0.11 *μ*m). Cells are then binned by cell length (x-axis) with 1 pixel (0.11 *μ*m) in each bin. The number of foci in each pixel is counted for all daughter cells (mother cells not counted to avoid any aging effects (Thiermann et al., 2023)), and then normalized by the number of cells at the same cell length. Thus, the color in each pixel on the fork plots shows the probability of foci appearing at that y-position in the cell in that cell length.

For single-cell initiation and termination data, we did manual annotation and inspection using two different methods. First, we used the results from foci identification as described previously in (Si et al., 2019). To ensure the first method was not biased by foci identification, our second method directly tracked foci from the subtracted fluorescence images on Napari (a new feature in Napari-mm3). We found that both methods produced similar distributions of the initiation mass for the wildtype, Δ2^ATP^, and Δ4, while the foci in Δ2^ADP^ was too messy to be tracked faithfully by the second method. We then adopted the foci tracking results from the first method as in Figure 2 and Figure 3. The cell length at initiations and terminations is recorded, and the initiation mass is defined as the cell length at initiation, divided by the number of origins (given the negligible change in cell width (Taheri-Araghi et al., 2015)).

Additional details of strain information and data analysis are described in Supplementary Information.

## Notes

### Competing Interest Statement

The authors have declared no competing interest.

### Summary of Updates

rewrote the paper based on our initiator-titration model v2.

## Bibliography

Balleza, E., Kim, J. M., and Cluzel, P. (2018). Systematic characterization of maturation time of fluorescent proteins in living cells. Nat. Methods, 15(1):47–51.

Berger, M. and Wolde, P. R. T. (2022). Robust replication initiation from coupled homeostatic mechanisms. Nat. Commun., 13 (1):6556.

Bhat, D., Hauf, S., Plessy, C., Yokobayashi, Y., and Pigolotti, S. (2023). Correction: Speed variations of bacterial replisomes. Elife, 12.

Campbell, J. L. and Kleckner, N. (1990). E. coli oric and the dnaa gene promoter are sequestered from dam methyltransferase following the passage of the chromosomal replication fork. Cell, 62(5):967–979.

Charbon, G., Bjørn, L., Mendoza-Chamizo, B., Frimodt-Møller, J., and Løbner-Olesen, A. (2014). Oxidative DNA damage is instrumental in hyperreplication stress-induced inviability of escherichia coli. Nucleic Acids Res., 42(21):13228–13241.

Charbon, G., Klitgaard, R. N., Liboriussen, C. D., Thulstrup, P. W., Maffioli, S. I., Donadio, S., and Løbner-Olesen, A. (2018). Iron chelation increases the tolerance of escherichia coli to hyper-replication stress. Sci. Rep., 8(1):1–14.

Charbon, G., Mendoza-Chamizo, B., Campion, C., Li, X., Jensen, P. R., Frimodt-Møller, J., and Løbner-Olesen, A. (2021). Energy starvation induces a cell cycle arrest in escherichia coli by triggering degradation of the DnaA initiator protein. Front Mol Biosci, 8:629953.

Cherepanov, P. P. and Wackernagel, W. (1995). Gene disruption in escherichia coli: TcR and KmR cassettes with the option of flp-catalyzed excision of the antibiotic-resistance determinant. Gene, 158(1):9–14.

Datsenko, K. A. and Wanner, B. L. (2000). One-step inactivation of chromosomal genes in Escherichia coli K-12 using PCR products. Proceedings of the National Academy of Sciences, 97(12):6640–6645.

Donachie, W. D. (1968). Relationship between cell size and time of initiation of DNA replication. Nature, 219(5158):1077–1079.

Donachie, W. D. and Blakely, G. W. (2003). Coupling the initiation of chromosome replication to cell size in escherichia coli. Curr. Opin. Microbiol., 6(2):146–150.

Erzberger, J. P., Mott, M. L., and Berger, J. M. (2006). Structural basis for ATP-dependent DnaA assembly and replication-origin remodeling. Nat. Struct. Mol. Biol., 13(8):676–683.

Frimodt-Møller, J., Charbon, G., Krogfelt, K. A., and Løbner-Olesen, A. (2015). Control regions for chromosome replication are conserved with respect to sequence and location among escherichia coli strains. Front. Microbiol., 6:1011.

Frimodt-Møller, J., Charbon, G., Krogfelt, K. A., and Løbner-Olesen, A. (2016). DNA replication control is linked to genomic positioning of control regions in escherichia coli. PLoS Genet., 12(9):e1006286.

Fu, H., Xiao, F., and Jun, S. (2023). Bacterial replication initiation as precision control by protein counting. PRX Life.

Fujimitsu, K., Senriuchi, T., and Katayama, T. (2009). Specific genomic sequences of e. coli promote replicational initiation by directly reactivating ADP-DnaA. Genes Dev., 23(10):1221–1233.

Gon, S., Camara, J. E., Klungsøyr, H. K., Crooke, E., Skarstad, K., and Beckwith, J. (2006). A novel regulatory mechanism couples deoxyribonucleotide synthesis and DNA replication in escherichia coli. EMBO J., 25(5):1137–1147.

Gras, K., Fange, D., and Elf, J. The escherichia coli chromosome moves to the replisome. (2023).

Guyer, M. S., Reed, R. R., Steitz, J. A., and Low, K. B. (1981). Identification of a sex-factor-affinity site in e. coli as gamma delta. Cold Spring Harb. Symp. Quant. Biol., 45 Pt 1:135–140.

Hansen, F. G., Christensen, B. B., and Atlung, T. (1991). The initiator titration model: computer simulation of chromosome and minichromosome control. Res. Microbiol., 142(2):161–167.

Huang, D., Johnson, A. E., Sim, B. S., Lo, T. W., Merrikh, H., and Wiggins, P. A. (2023). The in vivo measurement of replication fork velocity and pausing by lag-time analysis. Nat. Commun., 14(1):1762.

Hunt, T. and Murray, A. The cell cycle : an introduction. Oxford University Press, (1993).

Jun, S., Si, F., Pugatch, R., and Scott, M. (2018). Fundamental principles in bacterial physiology—history, recent progress, and the future with focus on cell size control: a review. Rep. Prog. Phys., 81(5):056601.

Kasho, K. and Katayama, T. (2013). DnaA binding locus datA promotes DnaA-ATP hydrolysis to enable cell cycle-coordinated replication initiation. Proceedings of the National Academy of Sciences, 110(3):936–941.

Kasho, K., Fujimitsu, K., Matoba, T., Oshima, T., and Katayama, T. (2014). Timely binding of IHF and fis to DARS2 regulates ATP–DnaA production and replication initiation. Nucleic Acids Res., 42(21):13134–13149.

Kato, J. and Katayama, T. (2001). Hda, a novel DnaA-related protein, regulates the replication cycle in escherichia coli. EMBO J., 20(15):4253–4262.

Kawakami, H., Keyamura, K., and Katayama, T. (2005). Formation of an ATP-DnaA-specific initiation complex requires DnaA arginine 285, a conserved motif in the AAA+ protein family. J. Biol. Chem., 280(29):27420–27430.

Kitagawa, R., Ozaki, T., Moriya, S., and Ogawa, T. (1998). Negative control of replication initiation by a novel chromosomal locus exhibiting exceptional affinity for escherichia coli DnaA protein. Genes Dev., 12(19):3032–3043.

Knöppel, A., Broström, O., Gras, K., Elf, J., and Fange, D. (2023). Regulatory elements coordinating initiation of chromosome replication to the Escherichia coli cell cycle. Proceedings of the National Academy of Sciences, 120(22):e2213795120.

Le Treut, G., Si, F., Li, D., and Jun, S. (2021). Quantitative examination of five stochastic cell-cycle and cell-size control models for escherichia coli and bacillus subtilis. Front. Microbiol., 12:721899.

Leonard, A. C. and Grimwade, J. E. (2011). Regulation of DnaA assembly and activity: taking directions from the genome. Annu. Rev. Microbiol., 65:19–35.

Løbner-Olesen, A., Skarstad, K., Hansen, F. G., von Meyenburg, K., and Boye, E. (1989). The DnaA protein determines the initiation mass of escherichia coli K-12. Cell, 57(5):881–889.

Løbner-Olesen, A., Marinus, M. G., and Hansen, F. G. (2003). Role of SeqA and dam in escherichia coli gene expression: a global/microarray analysis. Proc. Natl. Acad. Sci. U. S. A., 100(8):4672–4677.

Lu, M., Campbell, J. L., Boye, E., and Kleckner, N. (1994). SeqA: a negative modulator of replication initiation in e. coli. Cell, 77 (3):413–426.

Mangiameli, S. M., Cass, J. A., Merrikh, H., and Wiggins, P. A. (2018). The bacterial replisome has factory-like localization. Curr. Genet., 64(5):1029–1036.

McGarry, K. C., Ryan, V. T., Grimwade, J. E., and Leonard, A. C. (2004). Two discriminatory binding sites in the escherichia coli replication origin are required for DNA strand opening by initiator DnaA-ATP. Proc. Natl. Acad. Sci. U. S. A., 101(9):2811–2816.

Miller, J. H. Experiments in Molecular Genetics. Cold Spring Harbor Laboratory, (1972).

Nakamura, K. and Katayama, T. (2010). Novel essential residues of hda for interaction with DnaA in the regulatory inactivation of DnaA: unique roles for hda AAA box VI and VII motifs. Mol. Microbiol., 76(2):302–317.

Neidhardt, F. C., Bloch, P. L., and Smith, D. F. (1974). Culture medium for enterobacteria. J. Bacteriol., 119(3):736–747.

Nishida, S., Fujimitsu, K., Sekimizu, K., Ohmura, T., Ueda, T., and Katayama, T. (2002). A nucleotide switch in the escherichia coli DnaA protein initiates chromosomal replication: evidence from a mutant DnaA protein defective in regulatory ATP hydrolysis in vitro and in vivo. J. Biol. Chem., 277(17):14986–14995.

Olivi, L., Berger, M., Creyghton, R. N. P., De Franceschi, N., Dekker, C., Mulder, B. M., Claassens, N. J., ten Wolde, P. R., and van der Oost, J. (2021). Towards a synthetic cell cycle. Nat. Commun., 12(1):1–11.

Ozaki, S., Kawakami, H., Nakamura, K., Fujikawa, N., Kagawa, W., Park, S.-Y., Yokoyama, S., Kurumizaka, H., and Katayama, T. (2008). A common mechanism for the ATP-DnaA-dependent formation of open complexes at the replication origin. J. Biol. Chem., 283(13):8351–8362.

Pelletier, J. F., Sun, L., Wise, K. S., Assad-Garcia, N., Karas, B. J., Deerinck, T. J., Ellisman, M. H., Mershin, A., Gershenfeld, N., Chuang, R.-Y., Glass, J. I., and Strychalski, E. A. (2021). Genetic requirements for cell division in a genomically minimal cell. Cell, 184(9):2430–2440.e16.

Pflug, F., Bhat, D., and Pigolotti, S. Genome replication in asynchronously growing microbial populations. (2023).

Reyes-Lamothe, R., Sherratt, D. J., and Leake, M. C. (2010). Stoichiometry and architecture of active DNA replication machinery in escherichia coli. Science, 328(5977):498–501.

Riber, L., Olsson, J. A., Jensen, R. B., Skovgaard, O., Dasgupta, S., Marinus, M. G., and Løbner-Olesen, A. (2006). Hda-mediated inactivation of the DnaA protein and dnaa gene autoregulation act in concert to ensure homeostatic maintenance of the escherichia coli chromosome. Genes Dev., 20(15):2121–2134.

Riber, L., Frimodt-Møller, J., Charbon, G., and Løbner-Olesen, A. (2016). Multiple DNA binding proteins contribute to timing of chromosome replication in e. coli. Front Mol Biosci, 3:29.

Ronneberger, O., Fischer, P., and Brox, T. U-Net: Convolutional networks for biomedical image segmentation. In Medical Image Computing and Computer-Assisted Intervention – MICCAI 2015, pages 234–241. Springer International Publishing, (2015).

Sauls, J. T., Cox, S. E., Do, Q., Castillo, V., Ghulam-Jelani, Z., and Jun, S. (2019). Control of bacillus subtilis replication initiation during physiological transitions and perturbations. MBio, 10(6).

Sauls, J. T., Schroeder, J. W., Brown, S. D., Le Treut, G., Si, F., Li, D., Wang, J. D., and Jun, S. Mother machine image analysis with MM3. (2019).

Sekimizu, K., Bramhill, D., and Kornberg, A. (1987). ATP activates dnaa protein in initiating replication of plasmids bearing the origin of the e. coli chromosome. Cell, 50(2):259–265.

Si, F., Li, D., Cox, S. E., Sauls, J. T., Azizi, O., Sou, C., Schwartz, A. B., Erickstad, M. J., Jun, Y., Li, X., and Jun, S. (2017). Invariance of initiation mass and predictability of cell size in escherichia coli. Curr. Biol., 27(9):1278–1287.

Si, F., Le Treut, G., Sauls, J. T., Vadia, S., Levin, P. A., and Jun, S. (2019). Mechanistic origin of Cell-Size control and homeostasis in bacteria. Curr. Biol., 29(11):1760–1770.e7.

Sompayrac, L. and Maaløe, O. (1973). Autorepressor model for control of DNA replication. Nat. New Biol., 241(109):133–135.

Su’etsugu, M., Takata, M., Kubota, T., Matsuda, Y., and Katayama, T. (2004). Molecular mechanism of DNA replication-coupled inactivation of the initiator protein in escherichia coli: interaction of DnaA with the sliding clamp-loaded DNA and the sliding clamp-hda complex. Genes Cells, 9(6):509–522.

Su’etsugu, M., Shimuta, T.-R., Ishida, T., Kawakami, H., and Katayama, T. (2005). Protein associations in DnaA-ATP hydrolysis mediated by the hda-replicase clamp complex. J. Biol. Chem., 280(8):6528–6536.

Su’etsugu, M., Harada, Y., Keyamura, K., Matsunaga, C., Kasho, K., Abe, Y., Ueda, T., and Katayama, T. (2013). The DnaA n-terminal domain interacts with hda to facilitate replicase clamp-mediated inactivation of DnaA. Environ. Microbiol., 15(12): 3183–3195.

Sugiyama, R., Kasho, K., Miyoshi, K., Ozaki, S., Kagawa, W., Kurumizaka, H., and Katayama, T. (2019). A novel mode of DnaA–DnaA interaction promotes ADP dissociation for reactivation of replication initiation activity. Nucleic Acids Res., 47(21): 11209–11224.

Taheri-Araghi, S., Bradde, S., Sauls, J. T., Hill, N. S., Levin, P. A., Paulsson, J., Vergassola, M., and Jun, S. (2015). Cell-Size control and homeostasis in bacteria. Curr. Biol., 25(3):385–391.

Theisen, P. W., Grimwade, J. E., Leonard, A. C., Bogan, J. A., and Helmstetter, C. E. (1993). Correlation of gene transcription with the time of initiation of chromosome replication in escherichia coli. Mol. Microbiol., 10(3):575–584.

Thiermann, R., Sandler, M., Ahir, G., Sauls, J. T., Schroeder, J. W., Brown, S. D., Le Treut, G., Si, F., Li, D., Wang, J. D., and Jun, S. (2023). Tools and methods for high-throughput single-cell imaging with the mother machine. bioRxiv.

Thomason, L. C., Sawitzke, J. A., Li, X., Costantino, N., and Court, D. L. (2014). Recombineering: genetic engineering in bacteria using homologous recombination. Curr. Protoc. Mol. Biol., 106:1.16.1–1.16.39.

Wallden, M., Fange, D., Lundius, E. G., Baltekin, Ö., and Elf, J. (2016). The synchronization of replication and division cycles in individual e. coli cells. Cell, 166(3):729–739.

Wang, P., Robert, L., Pelletier, J., Dang, W. L., Taddei, F., Wright, A., and Jun, S. (2010). Robust growth of escherichia coli. Curr. Biol., 20(12):1099–1103.

Wold, S., Skarstad, K., Steen, H. B., Stokke, T., and Boye, E. (1994). The initiation mass for DNA replication in escherichia coli K-12 is dependent on growth rate. EMBO J., 13(9):2097–2102.

Youngren, B., Nielsen, H. J., Jun, S., and Austin, S. (2014). The multifork escherichia coli chromosome is a self-duplicating and self-segregating thermodynamic ring polymer. Genes Dev., 28(1):71–84.

Zheng, H., Bai, Y., Jiang, M., Tokuyasu, T. A., Huang, X., Zhong, F., Wu, Y., Fu, X., Kleckner, N., Hwa, T., and Liu, C. (2020). General quantitative relations linking cell growth and the cell cycle in escherichia coli. Nat Microbiol, 5(8):995–1001.

