## Supplementary material for "Robust control of replication initiation in the absence of DnaA-ATP ↔ DnaA-ADP regulatory elements in *Escherichia coli*": NA

### Supplementary Information

| Strain name | Alias | Genotype | Construction method | Usages |
|---|---|---|---|---|
| ALO8238 | wildtype | MG1655 Δ*lacIZYA Z1 (lacI tetR SpR)* | | flow cytometry experiments, Figure 3.1, 4.2, & S5 |
| ALO8112 | $\Delta 4$ | MG1655 Δ*lacIZYA* Δ*datA* Δ*hda::cmR* Δ*DARS1* Δ*DARS2 Z1(lacI tetR SpR)* | P1 transduction | flow cytometry experiments, Figure 3.1, 4.2, & S5 |
| ALO8337 | $\Delta 2^{\mathrm{ADP}}$ | MG1655 Δ*lacIZYA* Δ*datA* Δ*hda::cmR Z1(lacI tetR SpR)* | P1 transduction | flow cytometry experiments, Figure 2.1, & S5 |
| ALO8340 | $\Delta 2^{\mathrm{ATP}}$ | MG1655 Δ*lacIZYA* Δ*DARS1* Δ*DARS2 Z1(lacI tetR SpR)* | P1 transduction | flow cytometry experiments, Figure 2.1, & S5 |
| ALO7931 (SJ2206) | wildtype | ALO8238 *dnaA kanR yPet-dnaN* | P1 transduction | mother machine experiments, Figure 1.3, 2.1, 3.1, 3.2, S1, & S4.2 |
| ALO7918 (SJ2222) | $\Delta 4$ | ALO8112 *dnaA kanR yPet-dnaN* | P1 transduction | mother machine experiments, Figure 3.1, 3.3, 3.4, 4.3, 4.4, S1, & S4.3 |
| SJ2273 | $\Delta 2^{\mathrm{ADP}}$ | ALO8337 *dnaA kanR yPet-dnaN* | P1 transduction | mother machine experiments, Figure 2.1, 2.4, 2.5, & S1 |
| SJ2219 | $\Delta 2^{\mathrm{ATP}}$ | ALO8340 *dnaA kanR yPet-dnaN* | P1 transduction | mother machine experiments, Figure 2.1, 2.2, 2.3, & S1 |
| ALO8450 | wildtype DnaA$^{\mathrm{wt}}$ | ALO8238 *pLR40_Plac::dnaA*$^{\mathrm{wt}}$ | plasmid transformation | flow cytometry experiments, Figure 4.1 |
| ALO8451 | wildtype DnaA$^{\mathrm{R334A}}$ | ALO8238 *pLR40_Plac::dnaA*$^{\mathrm{R334A}}$ | plasmid transformation | flow cytometry experiments, Figure S2.1 |
| ALO8452 | wildtype DnaA$^{\mathrm{T174P}}$ | ALO8238 *pLR40_Plac::dnaA*$^{\mathrm{T174P}}$ | plasmid transformation | flow cytometry experiments, Figure S2.2 |
| ALO8446 | $\Delta 4$ DnaA$^{\mathrm{wt}}$ | ALO8112 *pLR40_Plac::dnaA*$^{\mathrm{wt}}$ | plasmid transformation | flow cytometry experiments, Figure 4.1 |
| ALO8447 | $\Delta 4$ DnaA$^{\mathrm{R334A}}$ | ALO8112 *pLR40_Plac::dnaA*$^{\mathrm{R334A}}$ | plasmid transformation | flow cytometry experiments, Figure S2.1 |
| ALO8448 | $\Delta 4$ DnaA$^{\mathrm{T174P}}$ | ALO8112 *pLR40_Plac::dnaA*$^{\mathrm{T174P}}$ | plasmid transformation | flow cytometry experiments, Figure S2.2 |
| SJ2060 | wildtype | ALO7931 *dnaA gfpmut2 ampR mcherry-dnaN* | $\lambda$ red recombineering | mother machine experiments, Figure 5.3 |
| SJ2061 | $\Delta 4$ | ALO7918 *dnaA gfpmut2 ampR mcherry-dnaN* | $\lambda$ red recombineering | mother machine experiments, Figure 5.3 |

**Table 1.** Strain information of *Escherichia coli* K-12.

| Strain | Genotype | Mutations found by sequencing |
|---|---|---|
| ALO7915 | *ΔlacIZYA ΔdatA Δhda ΔDARS1 ΔDARS2* | NC_000913.3 : Δ 3,555,471-3,558,888 (Δ malT rtcA rtcB rtcR), NC_000913.3: 371758 IS insertion (mhpC), NC_000913.3: 1879827 IS insertion (yeaR) |
| ALO7916 | *ΔlacIZYA ΔdatA Δhda ΔDARS1 ΔDARS2* | NC_000913.3 : Δ 3,555,471-3,558,888 (Δ malT rtcA rtcB rtcR), NC_000913.3: 371758 IS insertion (mhpC), NC_000913.3: 1879827 IS insertion (yeaR) |
| ALO8238 | *ΔlacIZYA Z1 (lacI tetR SpR)* | NC_000913.3 : 2,132,470 A→T (wcaB) |
| ALO8112 | *ΔlacIZYA ΔdatA Δhda ΔDARS1 ΔDARS2 Z1(lacI tetR SpR)* | NC_000913.3 : Δ 3,555,471-3,558,888 (Δ malT rtcA rtcB rtcR), NC_000913.3: 371758 IS insertion (mhpC), NC_000913.3: 1879827 IS insertion (yeaR) |
| ALO8337 | *ΔlacIZYA ΔdatA Δhda Z1(lacI tetR SpR)* | NC_000913.3 : 2,132,470 A→T (wcaB), NC_000913.3 : 803,662 C→A (ybhJ) |
| ALO8339 | *ΔlacIZYA ΔdatA Δhda Z1(lacI tetR SpR)* | NC_000913.3 : 2,132,470 A→T (wcaB), NC_000913.3 : 803,662 C→A (ybhJ) |
| ALO8340 | *ΔlacIZYA ΔDARS1 ΔDARS2 Z1(lacI tetR SpR)* | NC_000913.3 : 3,554,106 25bp insertion (malT), NC_000913.3: 371758 IS insertion (mhpC), NC_000913.3: 1879827 IS insertion (yeaR) |
| ALO8341 | *ΔlacIZYA ΔDARS1 ΔDARS2 Z1(lacI tetR SpR)* | NC_000913.3 : 803,662 C→A (ybhJ), 3,553,331 C→T (malT), NC_000913.3: 371758 IS insertion (mhpC), NC_000913.3: 1879827 IS insertion (yeaR) |

**Table 2.** Whole-genome sequencing results. The sequences have been uploaded on NCBI as a biosample. BioProject ID: PRJNA1034260.

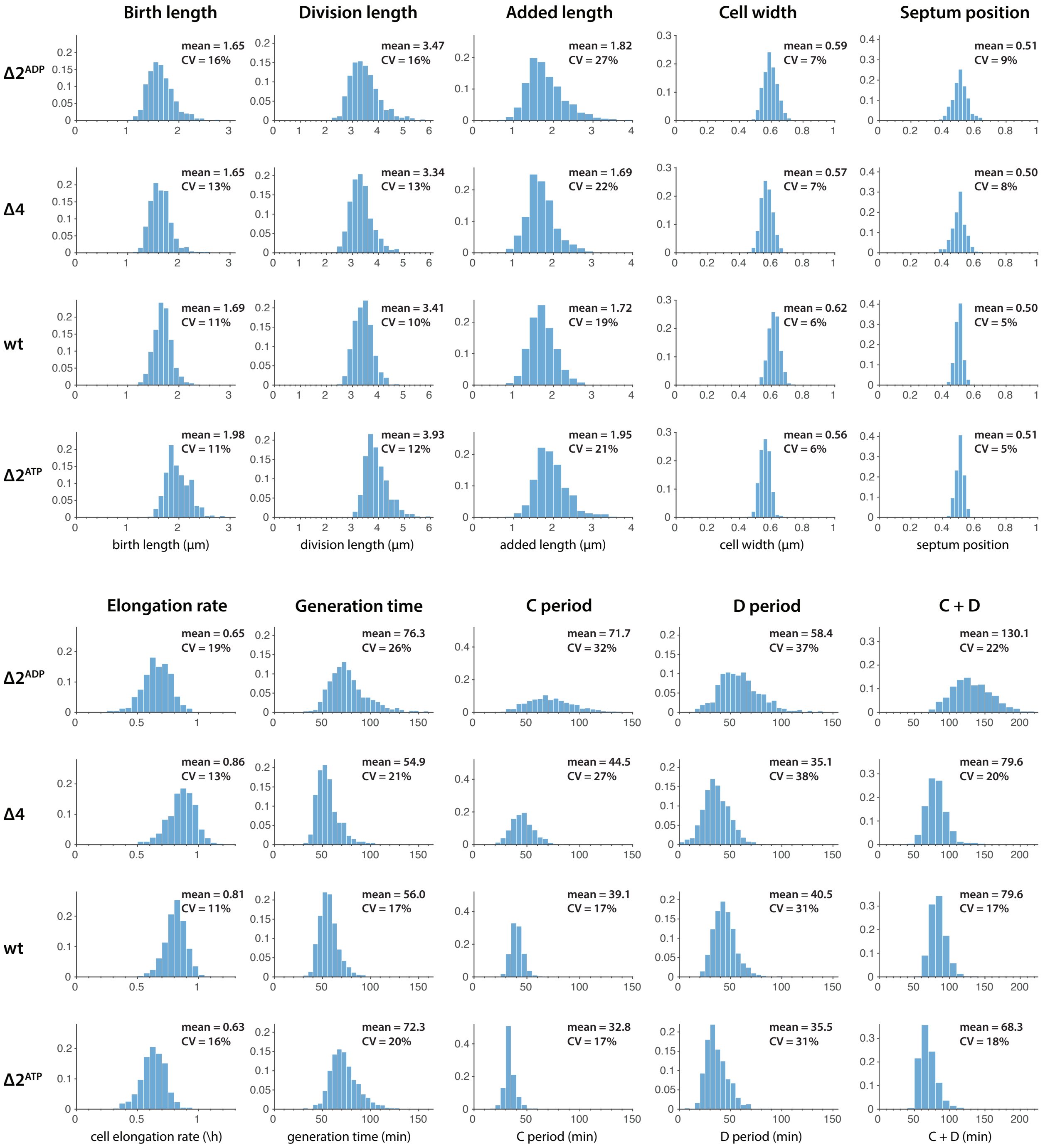


**Figure S1.** Distributions of physiological parameters from mother machine experiments. Only the second daughter cells are counted to avoid cell aging effects in mother cells. The y-axis shows the probability density. Number of cells counted in each strain: $\Delta 2^{\mathrm{ADP}}$: 617, $\Delta 4$: 593, wildtype: 1017, $\Delta 2^{\mathrm{ATP}}$: 666.

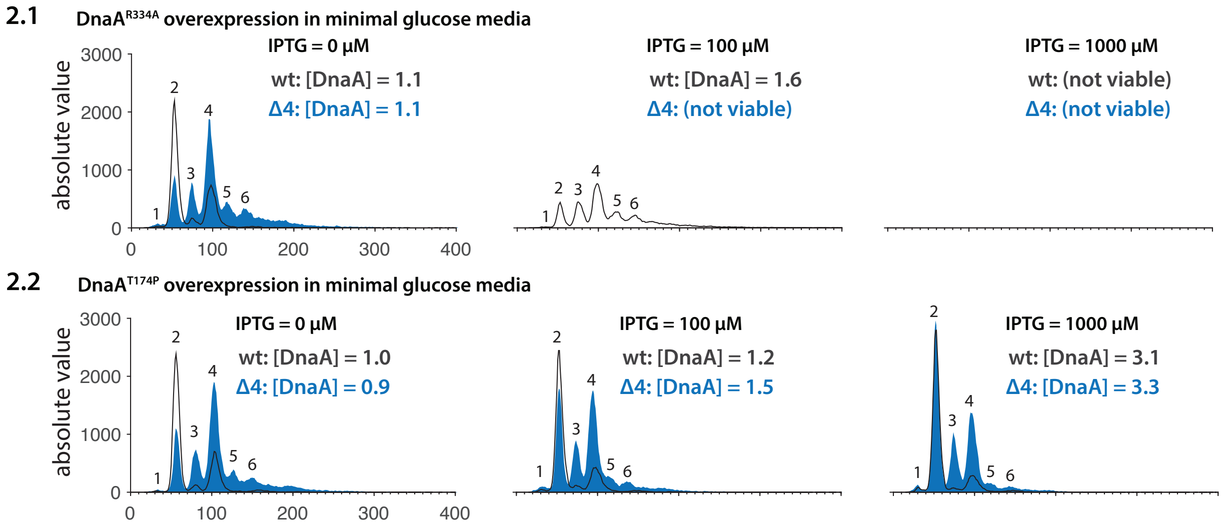


**Figure S2.** Flow cytometry experiments of DnaA mutant strains. **(2.1)** Expression of DnaA$^{\mathrm{R334A}}$ is induced by IPTG using a pLac promoter on a pLR40 plasmid in both wildetype and $\Delta 4$ *E. coli*. **(2.2)** Expression of DnaA$^{\mathrm{T174P}}$ is induced by IPTG using a pLac promoter on a pLR40 plasmid in both wildetype and $\Delta 4$ *E. coli*. The [DnaA] levels are shown in Figure S3.

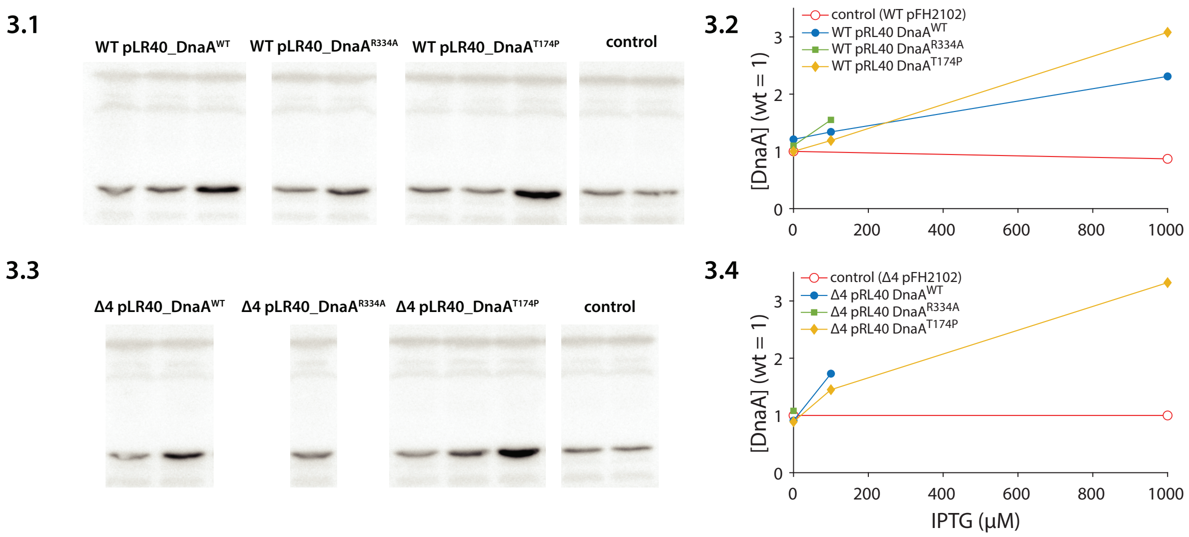


**Figure S3.** DnaA level measurements. **(3.1, 3.3)** Western blots of wildtype/$\Delta 4$ *E. coli* with different DnaA-mutant plasmids. The control plasmid has no $dnaA$ gene. **(3.2, 3.4)** DnaA induction curves of different strains.

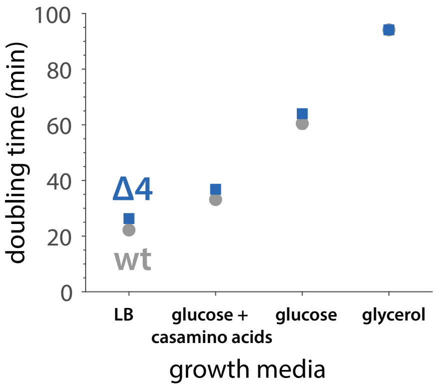


**Figure S4.** The average doubling time of wildtype and $\Delta 4$ *E. coli* in different growth conditions.

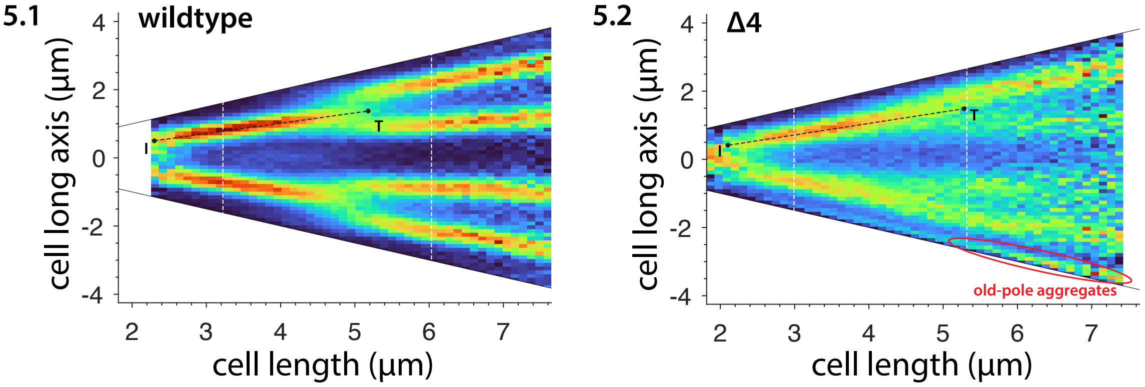


**Figure S5.** Fork plots of wildtype **(5.1)** and $\Delta 4$ **(5.2)** in MOPS rich glycerol (fast growth condition). The doubling time is 27 min (wt) vs. 35 min ($\Delta$4). In addition to the noisy foci distribution in $\Delta 4$ *E. coli*, $\Delta$4 also shows strong aging effects that quickly cause cell death or cell filamentation in 4 to 5 generations. As seen in (5.2), the old poles typically have aggregates in $\Delta$4 but not in wildtype, even though we only considered daughter cells, indicating the growth defect of $\Delta$4 may come from the old-pole aggregates. **Estimation of initiation mass, termination mass, and C period:** To estimate the initiation mass and the termination mass, we made two assumptions: 1. the distributions of the initiation mass (length) and the termination mass (length) are roughly symmetric (near-Gaussian distribution), and 2. the initiation mass distribution does not overlap with the termination mass distribution. Based on these two assumptions, we reason that if we track along a fork trajectory (black dashed lines), the initiation point (point I) and the termination point (point T) are when the maximal foci density drops by half. In (5.1), the wildtype cell length at point I and point T read $L_{\mathrm{I}} = 2.31$ $\mu$m and $L_{\mathrm{T}} = 5.17$ $\mu$m, respectively; in (5.2) the $\Delta 4$ cell length at point I and point T read $L_{\mathrm{I}} = 2.09$ $\mu$m and $L_{\mathrm{T}} = 5.28$ $\mu$m, respectively. By assuming exponential growth, we have $L_{\mathrm{T}} = L_{\mathrm{I}} 2^{\frac{C}{\tau}}$. Thus, we obtain $C/\tau \approx 1.2$ for wildtype, and $C/\tau \approx 1.3$ for $\Delta 4$.

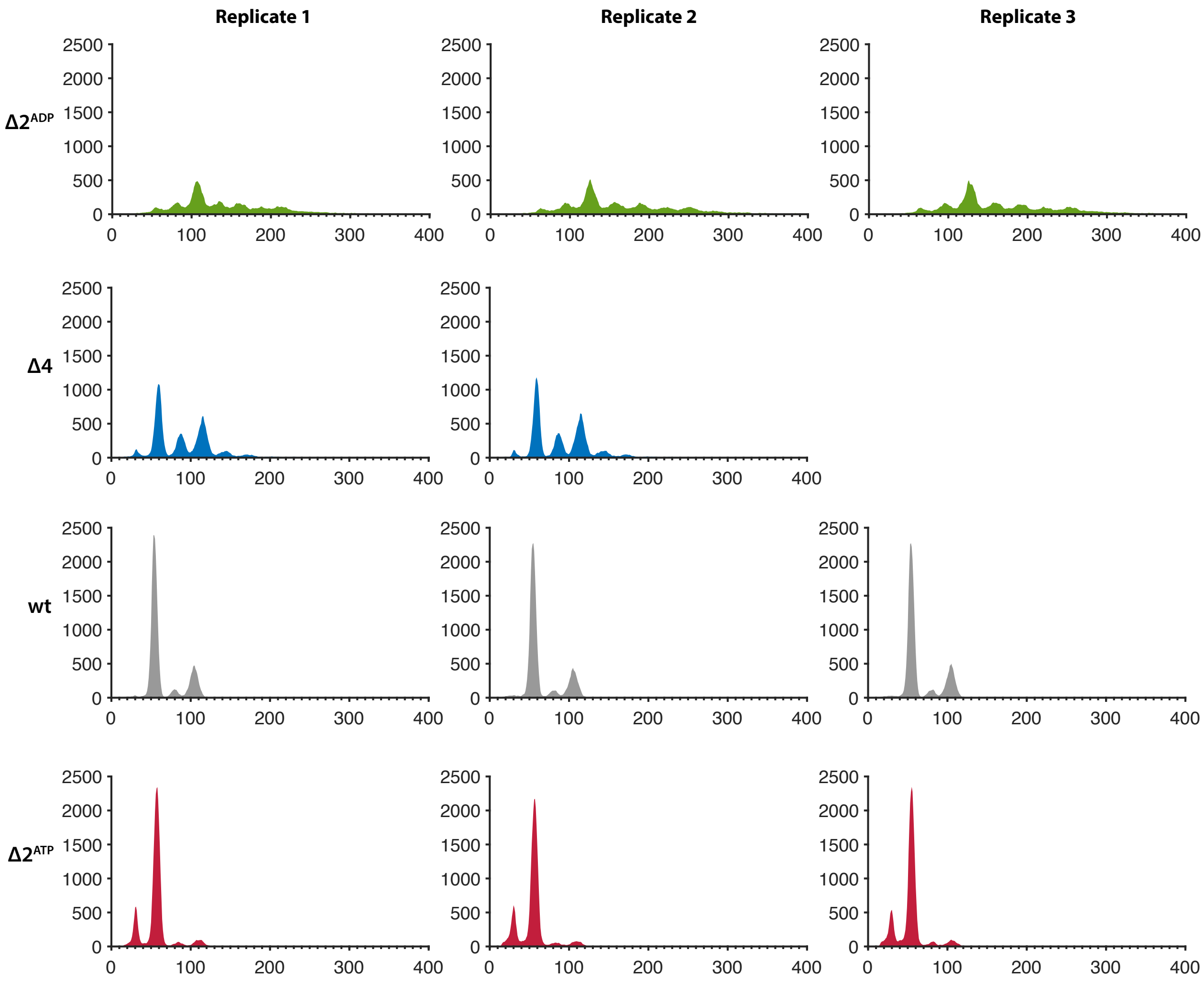


**Figure S6.** Replicates of flow cytometry experiments of $\Delta 2^{\mathrm{ADP}}$, $\Delta 4$, wildtype, and $\Delta 2^{\mathrm{ATP}}$ in glucose minimal media.

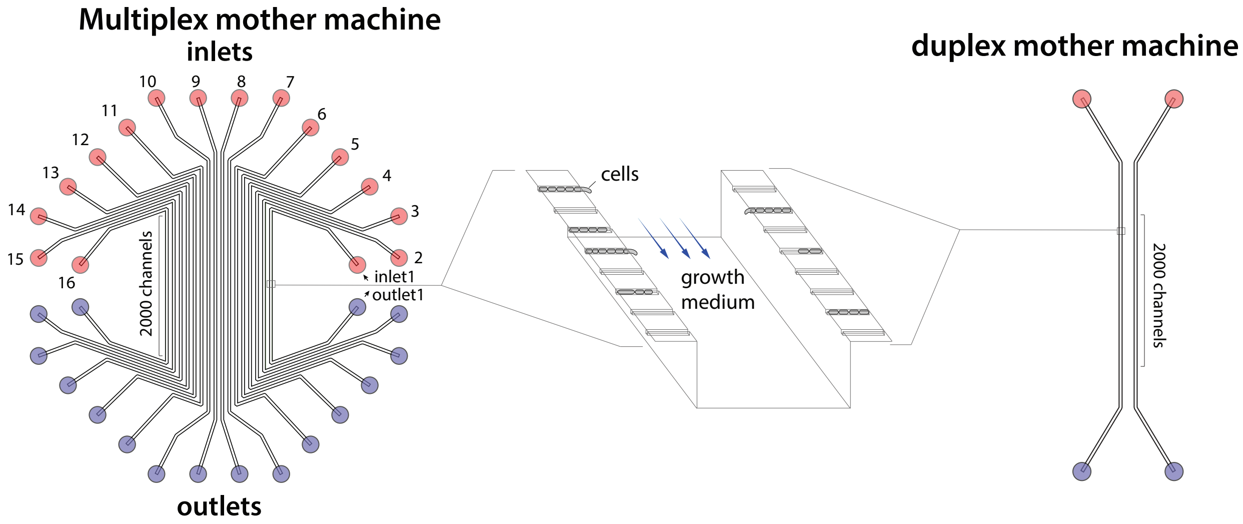


**Figure S7.** Designs of multiplex mother machines and duplex mother machines for parallel mother machine experiments.
